## Supporting Information PDF for "Synthetic community Hi-C benchmarking provides a baseline for virus-host inferences"

*Adsorption kinetics experiments*

Each phage–host pair was tested to determine which phages were able to adsorb to which host strains (**Table S3**). Each bacterial strain was inoculated and transferred as for the synthetic community sampling experiments. Once each strain had reached mid-exponential phase and at least $1{\times}10^8$ CFU/mL, adsorption experiments were performed for each phage–host pair in duplicate. Cells and phage were incubated together in 1 mL PC medium at a multiplicity of infection (MOI) of 1 ($1{\times}10^8$ CFU/mL and $1{\times}10^8$ PFU/mL). A no bacteria control was also performed with each phage to ensure phage titer did not decrease due to other factors (e.g., sticking to the tube). For phages with no known host in the synthetic community (phages φ14:2 and φSM) or known poor adsorption to mock community bacteria (phage φST), their hosts of isolation were included as a positive control. Total and free phage were sampled and plated to determine PFU/mL at 0 min and 60 min, or 30 min where the known latent period is less than 60 min (phages HM1 and φ46:1). Adsorption was determined by comparing the free phage concentration at 60 min or 30 min to that at 0 min. A t-test was performed and any result that showed a significant decrease in free phage over time was considered positive for adsorption for that phage–host pair.

Follow-up adsorption kinetics experiments (**Table S3**) were conducted on phage–host pairs showing significant adsorption in the initial experiment above. These experiments were performed in biological triplicates at MOI=0.1 to reduce the chances of multiple adsorptions to a single host cell, and sampled every few minutes for ~30 min, but otherwise similar to the above. Some experiments did not include a 15 min timepoint – for these, data from 16 min is given instead.

*Cell recovery experiment after freeze-thaw cycle*

Strains H71, H105, and 13-15 were inoculated in PZM, and strain 18 in MLB, and shaken overnight at 150 rpm at room temperature. Inoculations were transferred to fresh PC media and incubated stationary at room temperature. The optical density (OD) was monitored and graphed every 30-60 minutes until it reached $1{\times}10^8$ cells/mL. Each transfer was then diluted to $1{\times}10^7$ cells/mL in PC phage buffer. 1mL of each dilution was pipetted into cryotubes with different preservatives (DMSO, betaine, glycerol, and no preservative, as in the SynCom sampling). Sets with preservatives were flash frozen in liquid nitrogen and stored at -80°C. Colony-forming units (CFUs) were plated from the no-preservative set, remaining samples were centrifuged, pellets were resuspended in PBS, filtered, and the resuspended pellets and filtrates were frozen at -20°C. Finally, CFUs were counted (Figure S4).

*Free DNA test*

We used qPCR to determine the concentration of free bacterial genomic DNA after pelleting of each strain before and after freeze-thaw in various preservatives (**Figure S4**). Each strain was inoculated and transferred as for the synthetic community sampling experiments. Once each strain had reached mid-exponential phase and at least 1e8 CFU/mL, we performed the freeze-thaw experiment in various preservatives in triplicate. Each strain was diluted to 1e7 CFU/mL and preserved as follows: 1) no preservative, 2) 6.5% DMSO, 3) 0.5M betaine, or 4) 20% glycerol, as well as 5) a fresh (non-frozen), no preservative control. Sample sets 1–4 were flash frozen in liquid nitrogen and stored at -80 °C. Sample set 5 was centrifuged for 10 min at 10,000g at 4 °C to pellet cells and the supernatant discarded. The pellets were resuspended in 1 mL PBS, and 0.5 mL was 0.2 μm filtered to remove cells. Both the filtrate and remaining 0.5 mL of unfiltered pellet resuspension were stored at -20 °C. Sample sets 1–4 were thawed and treated as set 5. qPCR was performed on a 7500 Fast Dx Real-Time PCR System (Applied Biosystems) with PerfeCTa SYBR Green FastMix Reaction Mix with low ROX (QuantaBio) in 10µl reactions. Per reaction, we used 5 µl PerfeCTa master mix, 0.3 µl 10mM forward primer, 0.3 µl 10mM reverse primer, 3.4 µl nuclease-free water, and 1 µl template.

The target PSA H71 has a forward sequence of TTCTGATTCTGATGCGCGTG and a reverse sequence of CTTCTGATGGATTAGCGCCG, with an annealing temperature of 63°C. The target PSA H105 has a forward sequence of TGTATCGCCTGCTTCACCTA and a reverse sequence of GCAGAACTTCCTACTTCCAGC, also with an annealing temperature of 63°C. The target PSA 13-15 has a forward sequence of GAGTTTGTGTCGTTGGATCGT and a reverse sequence of CCCAACTAGTAAACCACCAATCA, with an annealing temperature of 54°C. Lastly, the target CBA 18 has a forward sequence of TTTTACGAGAACGCCATCTTTCCAC and a reverse sequence of TGATGTAAGAGGGTTGAGGGCT, with an annealing temperature of 62°C.

Reactions were performed in technical duplicates with a standard curve consisting of 6x 10-fold dilution series of known concentration of the target strain DNA, which was used to calculate target sequence copies/µl. Cycling conditions were as follows: polymerase activation for 5 minutes at 95 °C; 40 cycles of 20 sec at 95 °C, 15 sec at primer annealing temperature (see SX), and 24 sec at 72 °C, and a 65 °–95 °C melt curve. Results were used to compare the number of DNA copies in the filtrate (free DNA) to that in the unfiltered, resuspended pellets (total DNA). The results from the frozen samples (sets 1–4) were compared to the fresh samples (set 5) to determine how free DNA changed with freeze-thaw for each preservative.

*Area Under the Curve for SynCom-1, SynCom-2 and SynCom-3 replicates*

This dataset summarizes the Area Under the Curve (AUC) values for replicates of synthetic microbial communities (SynCom-1, SynCom-2 and SynCom-3). AUC is a quantitative measure often used to evaluate the overall response or performance of a system over time, such as microbial growth, activity, or signal intensity. Higher AUC values typically indicate stronger or more sustained responses. However, these data contain too few data points to have any meaningful interpretation.

| Sample | AUC |
|---|---|
| SynCom-1 replicate 1 | 0.9464286 |
| SynCom-1 replicate 2 | 0.9627329 |
| SynCom-1 replicate 3 | 0.8739496 |
| SynCom-2 replicate 1 | 0.8333333 |
| SynCom-2 replicate 2 | 0.8000000 |
| SynCom-2 replicate 3 | 1.0000000 |
| SynCom-3 replicate 1 | 0.8333333 |
| SynCom-3 replicate 2 | 1.0000000 |
